# Viral infection arrests coccolithophore calcification and nutrient consumption, and triggers shifts in organic stoichiometry

**DOI:** 10.1101/2023.07.11.548577

**Authors:** Tamar Dikstein, Gilad Antler, Andre Pellerin, Shlomit Sharoni, Miguel J. Frada

**Affiliations:** Department of Ecology, Evolution and Behavior, Silberman Institute of Life Sciences, The Hebrew University of Jerusalem, Jerusalem, Israel; The Interuniversity Institute for Marine Sciences in Eilat, Eilat, Israel; Department of Earth & Environmental Sciences, Ben-Gurion University of the Negev, Beersheba, Israel; Institut des sciences de la mer, Université du Québec à Rimouski and GEOTOP, Rimouski, Québec, Canada; Department of Earth, Atmospheric and Planetary Sciences, Massachusetts Institute of Technology, Cambridge, United States of America

## Abstract

Blooms of the coccolithophore *Emiliania huxleyi* are routinely infected by a specific lytic virus (EhV), which rapidly kills host cells triggering bloom termination and organic and inorganic carbon export. However, the impact of EhV on the dynamic of resource acquisition and cellular stoichiometry remains unknown, limiting the current understanding of the ecological and biogeochemical significance of *E. huxleyi* blooms. To tackle this knowledge gap, we used algal and EhV cultures to determine over the course of infections the dynamics of alkalinity, modulated by calcification, nitrate and phosphate consumption and organic matter stoichiometry. We found that within 24hr alkalinity concentration stabilized and nutrient uptake declined to background levels. In parallel, the stoichiometric ratio of carbon to nitrogen was about 15% higher and the nitrogen to phosphorus ratio was about 12% lower during infections relative to controls. These variations likely resulted from lipid accumulation required for viral replication and the differential retention of phosphorus-rich macromolecular pools in decaying cells, respectively. Finally, after host population decay a progressive enrichment in phosphorus relative to nitrogen and carbon was detected in the remaining cell lysates. We estimate that this stoichiometric shift post-infection was driven by the progressive accumulation of heterotrophic bacteria involved in the degradation of organic material. Viral-mediated cell remodeling and consequent shifts in biomass stoichiometry likely impacts the patterns of nutrient cycling and biological carbon pump efficiency during large-scale blooms in the oceans.

## 1 Introduction

Phytoplankton are a taxonomically diverse group of photosynthetic microorganisms that inhabit the sunlit layer of the oceans where they are responsible for about 45% of global primary production and for driving the biological carbon pump (Field et al. 1998). This latter process results in a net drawdown of atmospheric CO_2_ into the ocean’s interior, henceforth influencing global climate (Sarmiento and Bender 1994). Biogeochemical cycles of the major nutrients (nitrogen; N and phosphorus; P) and carbon (C) are interlinked (Falkowski et al. 2008; Hutchins et al. 2009). The ratio of N to P in seawater determines which nutrient limits biological productivity in the ocean, with cascading effects on the marine food web (Finkel et al. 2010a; Deutsch and Weber 2012; Sharoni and Halevy 2020). In addition, the ratio of carbon to the limiting nutrient in organic matter, determines the maximum amount of carbon that can be fixed and exported (Galbraith and Martiny 2015). By consequence, phytoplankton elemental stoichiometry is a main component regulating the efficiency of the biological carbon pump.

Viruses are ubiquitous regulators of microbial growth and evolution in the oceans (Fuhrman 1999; Wilhelm and Suttle 1999). Mortality caused by lytic viruses is estimated to be a major constrain to marine primary production, food-web dynamics and nutrient cycles in the oceans (Mojica et al. 2016; Zimmerman et al. 2020). However, to our knowledge little is known on the impact of viral infections on phytoplankton and particulate organic carbon stoichiometry. Quantifying the impact of viral infection on nutrient cycling in phytoplankton cells is critical to improve our understanding of marine ecosystem functioning, biogeochemical cycles and the potential feedback on the global climate. Here, we address the impact of viral infections in *Emiliania huxleyi*, a cosmopolitan unicellular calcifying photoautotrophic organism belonging to the coccolithophore clade and capable of forming large-scale blooms in the oceans.

Coccolithophores are an important group of marine phytoplankton estimated to contribute 1% to 10% of primary production and phytoplankton biomass, increasing up to 40% during bloom conditions (Poulton et al. 2007, 2013). As a characteristic, coccolithophores produce an outer covering of calcite plates termed coccoliths (Young et al. 1999; Monteiro et al. 2016). This attribute confers coccolithophores a unique functional role in pelagic ecosystems. The biosynthesis of coccoliths is an energy costly process involving the active transport of the calcification substrates, Ca^2+^ and HCO^−^_3_, dissolved in seawater into the cell where coccoliths are precipitated (Monteiro et al. 2016). In doing so, coccolithophores reduce both seawater total dissolved inorganic carbon (DIC) and alkalinity, which changes the equilibrium between the different DIC forms and reduces the potential for seawater CO_2_ uptake and to buffer for pH changes (Rost and Riebesell 2004). Moreover, the biological precipitation and export of calcite results in a net release of CO_2_ to the atmosphere, a mechanism referred to as the alkalinity pump (or carbonate counter pump), which counter acts the biological carbon pump. Nonetheless, the impact of any CO_2_ production during calcification may be offset through the ballasting of organic carbon by heavy coccoliths entangled in marine snow, which can increase the efficiency of the biological carbon pump (Klaas and Archer 2002; Rost and Riebesell 2004). The relative strength of the two pumps, represented by the so-called rain ratio, determines to a large extent the air/sea exchange fluxes of CO_2_ and the magnitude of oceanic carbon sequestration, with important effects on the biogeochemical cycling of carbon and climate (Rost and Riebesell 2004).

In this context, blooms of *E. huxleyi* bear a significant impact. These blooms are typically extensive and develop over several weeks in temperate and subpolar waters during late spring/early summer periods (Holligan et al. 1993; Tyrrell and Merico 2004; Winter et al. 2014). The underlying conditions for bloom formation are numerous and may vary between sites, thus a matter of debate (Tyrrell and Merico 2004; Lessard et al. 2005). However, bloom decline is linked to an important extent to lytic infections by a specific *E. huxleyi* virus (EhV) (Bratbak et al. 1993; Brussaard et al. 1996; Wilson et al. 2002; Lehahn et al. 2014), a large double-stranded DNA Coccolithovirus (Schroeder et al. 2002). At the cellular level, EhV infection leads to profound remodelling of the *E. huxleyi* metabolism to support viral proliferation. These include the activation and recruitment of the host’s oxidative stress (Bidle et al. 2007; Vardi et al. 2009; Sheyn et al. 2016), upregulation of autophagy, nuclear degradation (Schatz et al. 2014), and both enhance glycolytic fluxes and lipid biosynthesis (Rosenwasser et al. 2014). The latter include triacylglycerols (Rosenwasser et al. 2014; Malitsky et al. 2016) and specific virus-derived glycosphingolipids that are important structural components of EhV particles and chemical signals triggering host’s programmed cell death (Vardi et al. 2009; Rosenwasser et al. 2014). Additionally, EhV infection prompts coccolith shedding from the cell surface, changing the optical signature of infected populations (Jacquet et al. 2002; Johns et al. 2019), and stimulates the production of transparent exopolymer particles (TEP) that can form large aggregates entrapping cells and coccoliths and enhance the vertical export flux of both particulate organic and inorganic carbon (Vardi et al. 2012; Laber et al. 2018).

Although focus has been directed to viral-mediate changes in metabolic and structural properties of host *E. huxleyi* cells, the impact of EhV on nutrient uptake, calcification dynamics and ultimately on the basal elemental stoichiometric compositions (C:N:P) of the viral infected cells (i.e., virocells, (Rosenwasser et al. 2016)) remains to be tested. This limits our understanding of basic feature of the host-virus interplay between *E. huxleyi* and EhV and bloom turnover, and the impact of viral infection on carbon export. To tackle this knowledge-gap, we examined the dynamic of macronutrient (N and P) consumption and calcification during viral infections, as well as the C:N:P stoichiometric composition of infected populations in controlled culture conditions. Our study demonstrates the impact of virus on the regulation of nutrient incorporation and cellular composition of *E. huxleyi* virocells and provide key insights on ensuing ecological and biochemical consequences of large-scale viral infections during oceanic blooms.

## 2 Methods

### 2.1 Cell culturing and viral-infections

The xenic calcifying *E. huxleyi* strain RCC1216 (diploid) was used in all experiments. Cell cultures were set in seawater-based K/2 (minus ammonium) medium (Keller et al. 1987), at 18°C, 13h:11h light: dark cycle, respectively, 80 μmol quanta m^−2^ s^−1^ irradiance using a warm white LED system and gently air-bubbled through 0.2 μm filter. A 6 L stock culture was grown for 3 days and then subdivided into six replicate glass bottles of 1 L each. Three of the replicates were infected with EhV 201 at a Multiplicity of infection (MOI) of 0.2, and three other replicates were used as non-infected controls. Sub-samples for cell, bacterial and viral counts, nitrate, phosphate, and alkalinity, as well as particulate organic carbon (POC), particulate organic nitrogen (PON), and particulate organic phosphorus (POP), were collected at different time points with high frequency in the first 72hr post-infection. An additional experiment run over 13 days was used to determine long term changes in POC, PON, and POP. See the details for sampling and analyses below.

### 2.2 Cell, virus, and bacterial counts

*E. huxleyi*, bacteria, and EhV particles were enumerated by flow cytometry (Attune NxT, Life Technology). Algal cells were identified based on chlorophyll fluorescence (excitation 488 nm) and side scatter (Marie et al. 1999; Frada et al. 2017). The mean side scatter and chlorophyll autofluorescence values were used as indicators for the degree of cell calcification (coccolith covering) and cell health, respectively. For viral and bacterial counts, samples of 0.5 mL were fixed with 0.5% glutaraldehyde and counterstained with 1:10000 SYBR Green (Invitrogen). Viruses and bacteria were identified by flow cytometry based on the green fluorescence (excitation: 488 nm and emission: 530–737 nm) and side scatter (Marie et al. 1999). The cell growth rate was determined as the natural logarithm of the cell density at 0 and 24hr divided by time. Viral burst size, was determined as the number of viruses released per lysed cell (estimated as the ratio of the maximal number of viruses produced to the maximum cell concentration reached by the specific host before cell decrease) (Bratbak et al. 1993; Jacquet et al. 2002).

### 2.3 Inorganic Nutrients and alkalinity

Duplicate samples of 10 mL were filtered through 0.2 μm filters (Sterivex, Millipore) and kept at 4°C until further analysis. Alkalinity was measured by acid-base titration following (Neder et al. 2022), via automatic titrator (Titroline 7000 Si analytics), and calculated using a gran plot with a precision lower than 20 mEq. Phosphate (as dissolved reactive phosphate) was measured by colorimetric spectroscopy using the Mo-Blue method (Cary 8454 UV-vis spectrometer, Agilent) with a precision of 0.1 mM and detection limit of 0.1 μM (Holman 1943). Nitrate was measured by second-derivative spectroscopy (Cary 8454 UV-vis spectrometer, Agilent) with a precision of 0.1 mM and detection limit of 0.1 μM (Crumpton et al. 1992).

### 2.4 Particulate organic carbon, nitrogen, and phosphorus

For POC and PON, 20 mL duplicate samples were collected and filtered under low-pressure vacuum filtration (100 mm Hg) onto pre-combusted (450°C, 5hr) Whatman GF/F filters (25 mm diameter, 0.7 μm effective pore size). Then the filters were rinsed with 230 μL of 1M hydrochloric acid (HCl) to remove particulate inorganic carbon (PIC), oven-dried overnight at 65°C, and folded into tin capsules. Carbon and nitrogen content analyses were conducted in Continuous-flow Isotope Ratio Mass Spectrometry (CF-IRMS) using a Deltaplus XP mass spectrometer (ThermoScientific) coupled to an elemental analysis (EA) COSTECH 4010 (Costech Analytical) equipped with a zero blank autosampler. For every 36 samples or less, 10 secondary standards are used to correct the data using a linear correction. Caffeine, nannochlopsis, and Mueller Hinton broth were used as secondary standards calibrated from NIST primary standard. Analytical error (n=50) on measurement was 0.06‰ and 0.08‰ for 6N and 6C, respectively. System suitability prior to analysis was evaluated using the standard deviation of zero reference gas (N and carbon dioxide) over 10 measurements and the maximum acceptable variation was set to 0.06%. Certified reference material from Elemental Microanalysis treated as a sample is used as quality control. Quantification is based on external secondary standards with a calibration range of 0.025 mg to 0.250 mg and 0.040 mg to 0.400 mg for nitrogen and carbon respectively according to (Agnihotri et al., 2014). For POP, 50 mL duplicate samples were filtered under low-pressure vacuum filtration (100 mm Hg) onto pre-combusted (450°C, 5hr) Whatman GF/F filters (25 mm diameter, 0.7 μm effective pore size). They were then combusted at 450°C, for 5hr in a muffle furnace, cooled down, and immersed in 10 ml of 0.5N of HCl for 60 min at 90°C in a heating block (Karl et al. 1991). Finally, 5 ml of supernatant was volumetrically subsampled into 15 ml centrifuge tubes for colorimetric spectroscopy using the Mo-Blue method (Cary 8454 UV-vis spectrometer, Agilent) as mentioned earlier for the analysis of inorganic P.

### 2.4 Mathematical estimation of bacterial contribution to particulate organic matter and statistical analyses

Given the rampant accumulation of free bacteria during the infection experiments (see Fig. 3), it is most likely that there was a concomitant accumulation of bacteria attached to particulate organic matter (Vincent et al. 2023). POM-bound bacteria were not measured. Thus, we estimated the contribution of bacteria to total particulate organic matter over the course of the experiment using a stoichiometric approach. These calculations were initialized 48hr post-infection, which coincided with the time of viral burst and increment in bacterial densities relative to control cultures. To calculate the fraction of bacteria out of the total particulate organic matter we used the following equation:

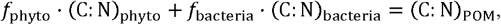

where (C: N)_phyto_, (C: N)_bacteria_, and (C: N)_POM_ are the C:N ratio in phytoplankton, bacteria, and POM, respectively. f_phyto_, and f_bacteria_ are the fraction of phytoplankton and bacteria, respectively. Assuming that f_phyto_ + f_bacteria_ =1, and rearranging, we get the following equation:

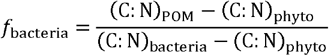

Here, (C: N)_phyto_= 8.32 is the ratio measured 48 hr post-infection (see Fig. 3 in the Results section). (C: N)_bacteria_= 4.94 is the average of elemental ratio of heterotrophic bacteria (Table S1 in Supplement S1); (Zimmerman et al. 2014; White et al. 2019) The (C: N)_POM_ is based on the results of our experimental findings (Fig. 5).

Additional comparisons between control and infected cultures were undertaken using student’s t-test with the Phyton (3.11.3).

## 3 Results

### 3.1 Algal growth and variations in optical properties

Detailed survey of viral infections was monitored during 72hr (Fig. 1A). During the first 36hr, cell growth was comparable between infections and controls, entailing a clear rise in cell density over the night period. After that, a clear cell decline was detected in the infections, while in the control cell density reached the stationary growth phase at ∼ 2.0 x 10^6^ cells mL^−1^ (Fig. 1A). During the second day, prior to cell decline, concurrent exponential accumulation of virions was detected in the infected cultures. Peak viral densities of ∼ 2.7 x 10^8^ virions mL^−1^ were detected 48hr post-infection. After that, viral standing stocks remaining relatively stable. The estimated viral burst size was estimated to be ∼230 viral particles cell^−1^.

**Figure 1.**
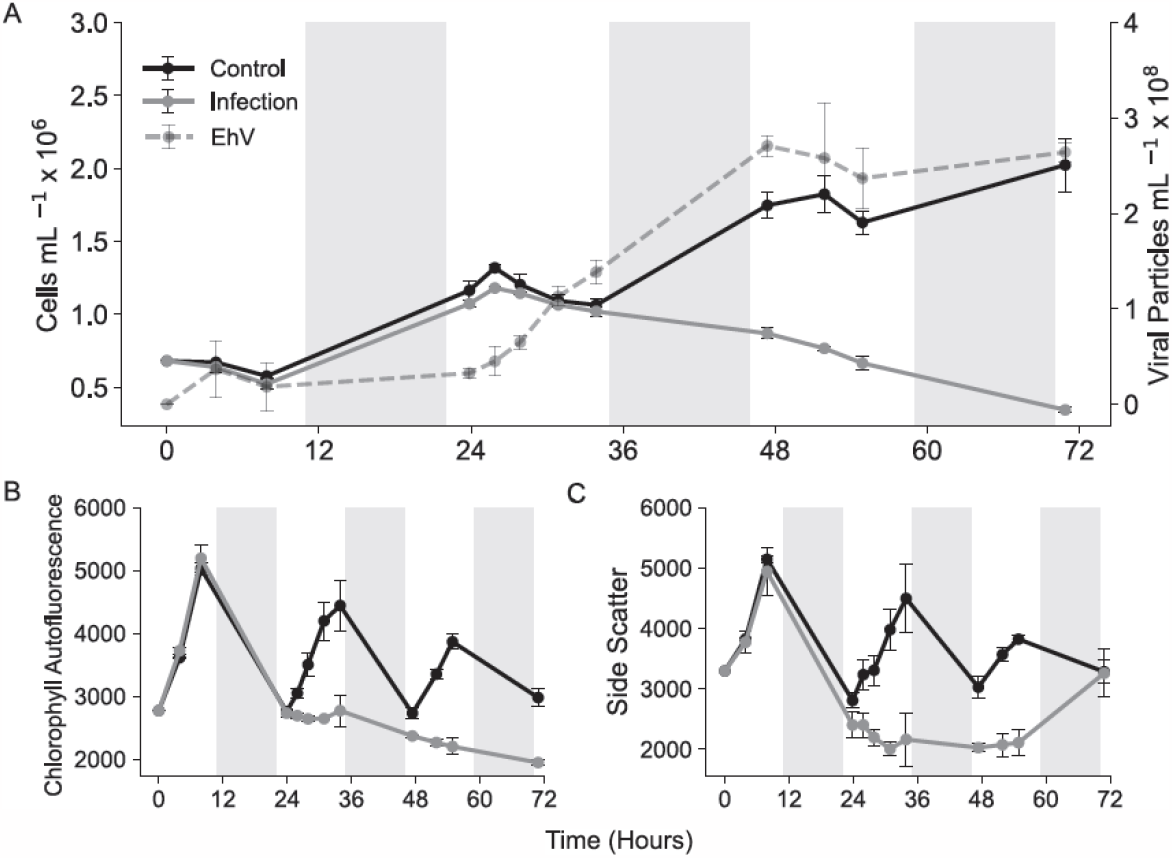
Algal and viral growth and variations in chlorophyll autofluorescence and side scatter. *E. huxleyi* cultures were infected with EhV 201 and compared with noninfected control cells. (A) Cell growth and extracellular viral production. (B) Mean population values of Red autofluorescence. (C) Mean population values of side scatter. The x-axes correspond to hours post infection. Grey sections represent the night period and dots indicate mean values of each treatment (Control/Infection), Error bars represent standard error of replicates, n=3.

Average chlorophyll autofluorescence and side scatter measured by flow cytometry were used to as indicators of cell health and degree of coccosphere integrity. During exponential growth, both parameters commonly oscillate through day-night cycles due to cell enlargement and coccolith buildup during exponential growth (Müller et al. 2008). This was detected during the first day in both treatments. However, during the second day, such diel oscillation in optical signals stopped and overall declined in infected cultures (Fig. 1B–C). A slight increase in the average side scatter signal was detected at 72hr. This can be explained by the formation of aggregates of decaying cellular materials with high scattering properties, not from the re-growth of remaining living cells.

### 3.2 Alkalinity, nitrate and phosphate consumption

During the first day, alkalinity (Fig. 2A), nitrate (Fig. 2B) and phosphate (Fig. 2C) draw down (i.e., due to coccolithophore consumption) was broadly similar in infected and control cultures. During the first light phase alkalinity, nitrate and phosphate removal (or consumption) averaged 13.40 ± 0.90 μmol L^−1^ h^−1^, 3.1 ± 0.4 μmol L^−1^ h^−1^ and 0.12 ± 0.06 μmol L^−1^ h^− 1^, respectively. During the first dark period alkalinity consumption was much lower (avg. 1.6 ± 0.8 μmol L^−1^ h^−1^), nitrate was about half (avg. 1.20 ± 0.20 μmol L^−1^ h^−1^), and phosphate similar to the light period (avg. 0.11 ± 0.01 μmol L^−1^). However, between 24hr and 36hr post-infection coincident with the arrest of cellular optical signals, alkalinity and nitrate consumption were both halted and phosphate consumption declined to residual levels (avg. 0.06 ± 0.04 μmol L^−1^) (Fig. 2A). After 36hr, phosphate removal stopped.

**Figure 2.**
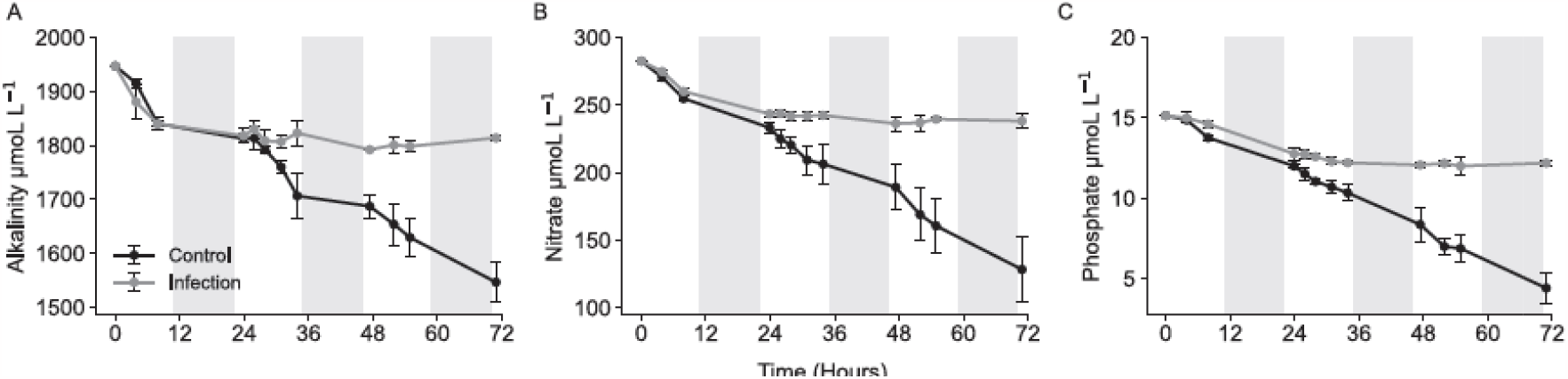
Dynamic of Alkalinity, nitrate, and phosphate in control and infected *E. huxleyi* cultures. The measured concentration of control and EhV infected cultures; (A) Alkalinity. (B) Dissolved nitrate. (C) Dissolved phosphate. The x-axes correspond to hours post-infection. Grey sections represent the night period and dots indicate the mean values of each treatment (Control/Infection), Error bars represent the standard error of replicates, n=3.

### 3.3 Particulate organic matter and stoichiometry dynamics

To determine the effect of viral infection on the elemental composition of coccolithophore cells, POC, PON, and POP were quantified over the length of the infection (72hr). An additional experiment, was set to determine the longer-term variations in the elemental composition of cell-derived lysates over 13 days post-infection (336hr). Both experiments were set up at the exact conditions and the output was nearly identical, thus presented together (unless otherwise indicated). POC, PON, and POP per cell as determined at the start of the experiment were 5.45 ± 0.15 pg cell^−1^, 1.22 ± 0.06 pg cell^−1^ and 0.14 ± 0.003 pg cell^−1^, respectively. We note that EhV infection leads to the rapid accumulation of cells in different physiological states, dead cells (Rosenwasser et al. 2014), TEP (Vardi et al. 2012), and aggregates (Vincent et al. 2021), all of which bear a particulate organic matter (POM) signature. Thus, the results for POC, PON, and POP are presented per unit volume to illustrate the temporal variations in the ensemble of particulate components accumulated in the cultures (Fig.3). The stoichiometric ratios are presented in Fig. 4, focusing separately on the infection period and post-infection for better visualization of variations during infections.

Over the infection period (72hr), the variations in cell and viral densities were broadly identical to Fig. 1A (Fig. 3A). Briefly, by 96hr cell densities in infected cultures were <10^3^ cell mL^−1^, while viral densities remained relatively stable until day 13. The control populations multiplied exponentially over the first 120hr and then transitioned to stationary growth phase. We note that in parallel, cohabiting bacteria progressively accumulated in the cultures (Fig. 3B). The increase in the bacterial population was higher in infected cultures doubling the density of controls after 96hr post-infection.

**Figure 3.**
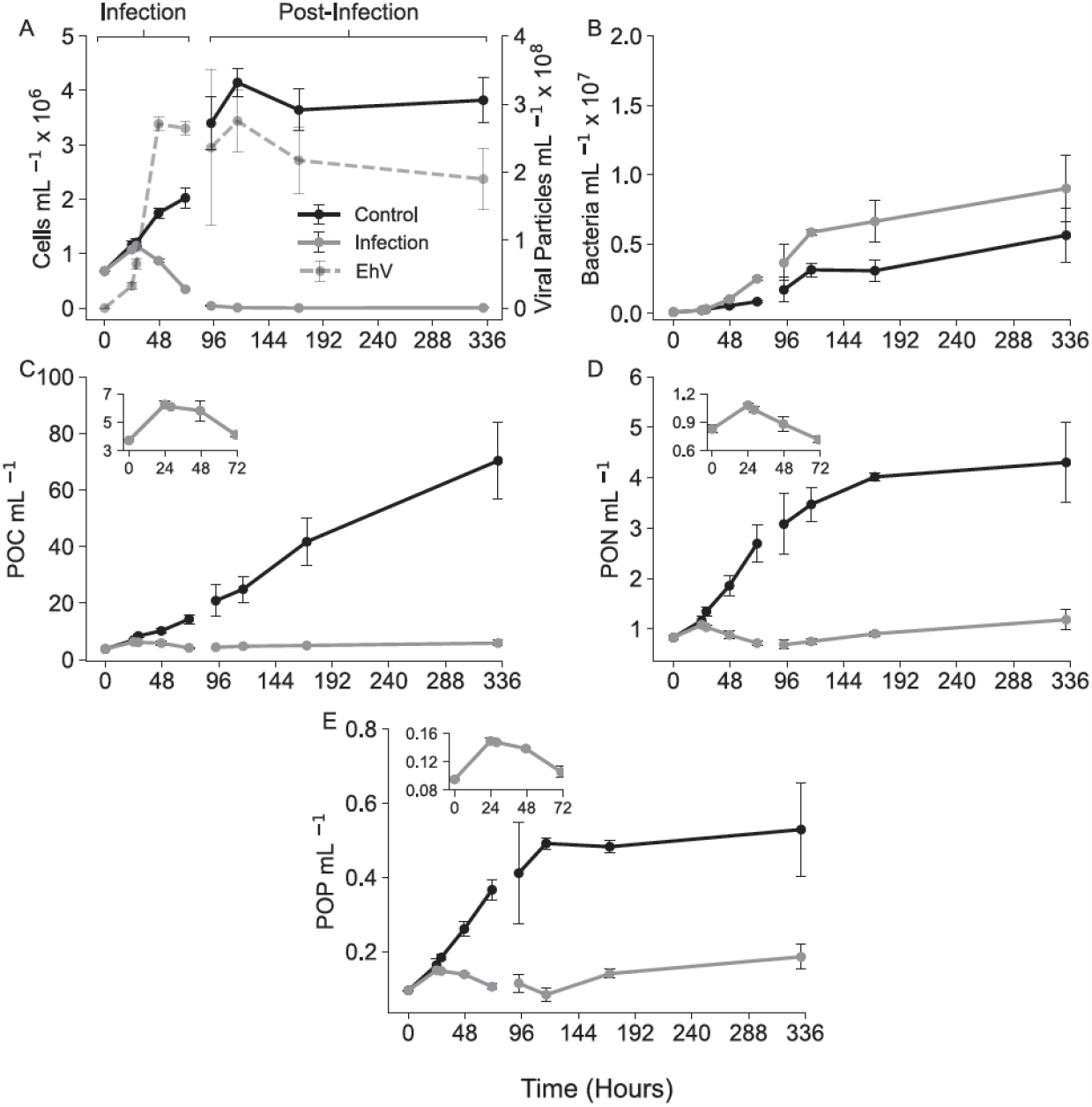
Variations in particulate organic matter (POC, PON, POP) in control and EhV-infected *E. huxleyi* cultures. Between 0hr and 72hr concerns the infection period. After 96hr concerns the post-infection period (see the main text for details). (A) Cell and viral densities. (B) Free bacterial densities (C–E) POC, PON, and POP concentrations, respectively. For higher detail, inset figures overlapping C, D and E highlight the variations in POC, PON, and POP during the infection period. Error bars represent the standard error of replicates, n=3.

POC, PON, and POP concentrations in the control cultures, progressively increased during the exponential phase as biomass accumulated (Fig. 3C-E). However, PON and POP stabilized as cultures entered stationary growth. In parallel, the molar stoichiometric ratios remained relatively stable during exponential growth and then C:N and C:P both increase during stationary growth (Fig. 4). By contrast, POC, PON, and POP concentrations in the presence of EhV increased during the first day of infection, but then rapidly declined paralleling cell demise. During the infection period, variations in the elemental stoichiometry relative to the controls were identified. Namely, the C:N ratio was significantly higher (C:N > 8.0) in infected cultures at 48hr and 72hr post-infection than in the controls (C:N ∼ 6.7) (Fig. 4A). By contrast, over the same period the average N:P was lower in infected cultures (N:P 14.0–15.5) than in the controls (N:P 15.7–16.2) (Fig. 4E). In the post-infection period, a slow progressive recovery in POC, PON, and POP concentrations was detected (Fig. 4B, D and F). However, in parallel, negative trends were detected for the C:N ratio (R^2^ = 0.91) and C:P ratio (R^2^ = 0.37), indicating a progressive enrichment of N and P relative to carbon in organic biomass post-infection.

**Figure 4.**
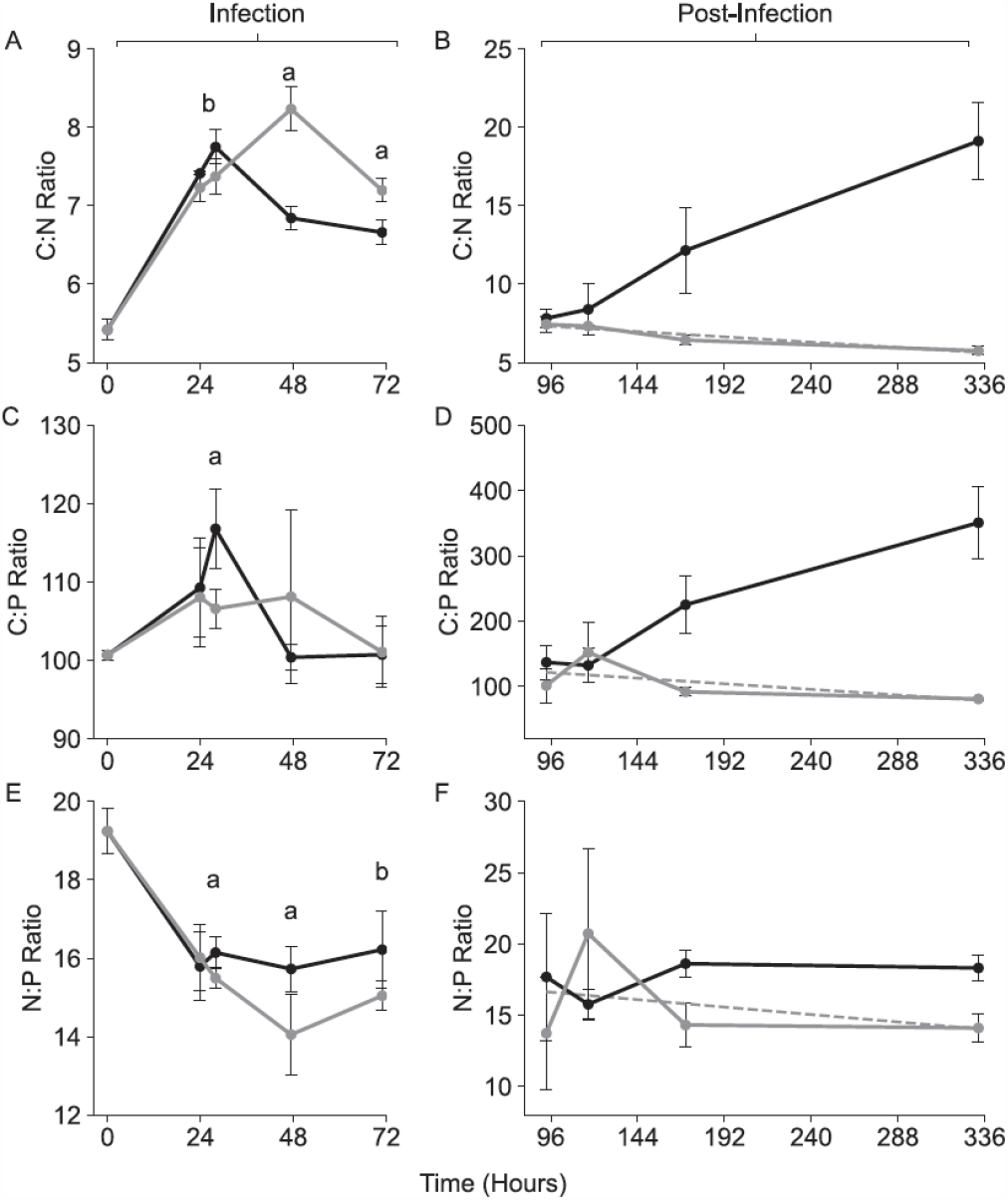
Variations in the elemental stoichiometric ratios (C:N, C:P, and N:P) in organic matter from control and EhV-infected *E. huxleyi* cultures. The infection period (0-72hr; A, C, and E) and post-infection period (from 96hr to 333hr; B, D, and F) are presented separately for better visualization. Respectively, the equations and R^2^ values of the linear regressions during the post-infection period for C:N, C:P and N:P are: y = -0.16x + 7.98, R^2^ = 0.91; y = - 4.28x + 138.39, R^2^ = 0.37; y = –0.26x + 17.65, R^2^ = 0.12. Error bars represent the standard error of replicates, n=3. Statistical significance is indicated by the superscript letter a (t-test, p < 0.05) and b (t-test, p < 0.1).

Bacterial stoichiometry often deviates from the typical phytoplankton Redfield Ratio. Average bacterial C:P is 77 and C:N is 4.9, while N:P is 17, similar to phytoplankton (Zimmerman et al. 2014; White et al. 2019). Thus, to test whether the observed shift in POM stoichiometry post-infection could result from the recycling of algal to bacterial biomass and accumulation of high bacterial titers, we calculated the contribution of bacteria over the course of our experiment. The results indicate that by the end of the infection at 72 hr post-infection, bacteria biomass contributed already around 30%, at 168hr (day 7) nearly 60%, and by day 13 nearly 80% of the total measured POM (Fig. 5).

**Figure 5.**
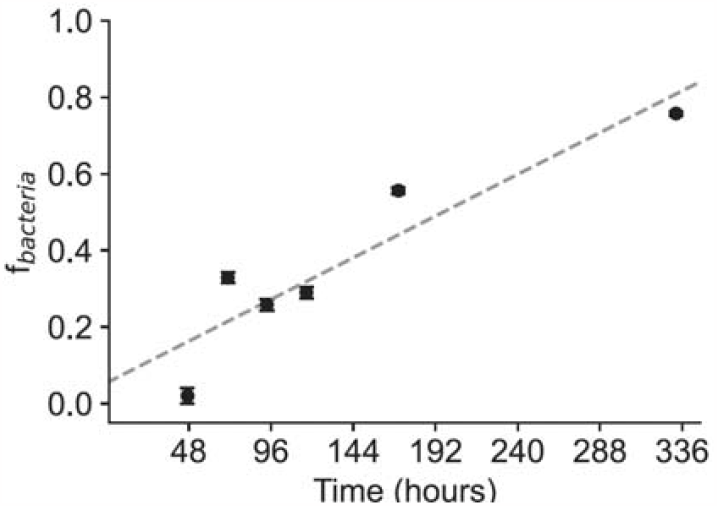
Mathematical estimates of the contribution of bacteria to particulate organic matter over the course of the infection and post-infection periods as defined in Fig. 3 and 4. The quadratic fitting describes the rate of bacterial accumulation, which is a saturation function (cannot exceed 1; y = 0.054x + 0.054, R^2^=0.85).

## 4 Discussion

### 4.1 EhV halts calcification and nutrient uptake in infected host cells

A typical dynamic of *E. huxleyi* (RCC 1216) by the specific EhV 201 set at an MOI of 0.2 was detected (Frada et al. 2008, 2017). Broadly, during the second day of infection, the daily oscillations of Chl fluorescence and side scattering signals were both halted and gradually declined in intensity (Fig. 1). These responses are consistent with the decline in the cellular chlorophyll content, impaired photophysiology (Evans et al. 2006; Kegel et al. 2007; Llewellyn et al. 2007; Kimmance et al. 2014), and coccolith shedding triggered during EhV infections (Johns et al. 2019). Then, after 36hr, the cell growth abruptly declined as a consequence of EhV-mediated cell death (Bidle et al. 2007; Vardi et al. 2009). A primary goal of this study, was to test the impact of EhV infection on 1) calcification as detected by determining the draw-down of alkalinity levels in the medium and 2) on the intake of the major macronutrients (nitrate and phosphate). During the first day, calcification was comparable between infected and control cultures. Alkalinity draw down occurred during the day, but not through the night (Fig. 2A). However, on the second day of infection, calcification was completely arrested. This coincided with the onset of viral burst and the overall shift in Chl fluorescence and side scatter signals in infected populations (Fig. 1). Coccolith biosynthesis is a complex energy-demanding mechanism requiring abundant lipid membranes for the coccolith vesicles and the secretion system that ultimately results in large inorganic structures seemingly unnecessary to viral replication (Monteiro et al. 2016). Thus, it is plausible that the viral-mediated breakdown of photosynthesis and the rapid progression of EhV, requiring large fluxes of nucleotides, lipidic resources, and energy, naturally results in the cessation of calcification and the incapability of coccolithophore virocells to keep and restore the disintegrating coccosphere.

As for calcification, nitrate and phosphate were both consumed during the first day in both control and infected cultures (Fig. 2B–C). However, in contrast to calcification, nutrient consumption continued through the dark period. Dark uptake of nitrate is known for *E. huxleyi* (Müller et al. 2008). Reports on the dark uptake of phosphate are less clear (Riegman et al. 2000; Müller et al. 2008). However, a higher affinity for phosphate has been described during the night, likely due to higher requirements for cell division (Geider and La Roche 2002). Nutrients acquired up to this stage through the infection are naturally incorporated into EhV biosynthesis as has been recently demonstrated in isotopic labeling experiments with carbon and N (Pasulka et al. 2018; Yakubovskaya et al. 2021). The reliance of EhV on external phosphate has not been thus far assessed, but likely follows analogous trends. During the second day nitrate consumption stopped. However, residual levels of phosphate removal were still detected through the day, which may be facilitated by the activity of a viral phosphate permease gene encoded by some EhV strains to sustain high viral demands for phosphorous (Wilson et al. 2005; Allen et al. 2006; Jover et al 2014). This occurred as side scatter and chlorophyll signals stopped oscillating, but prior and during initial stages of viral burst, which indicates that at this stage the virocell reached the final stages of viral maturation. Thus, from this moment further metabolic and energetic requirements of infected cells likely relied solely on intracellular recycling of available pool of energy and biomass resources.

### 4.2 EhV-mediated shifts in biomass stoichiometry during infection and post-infection

As expected, POC, PON, and POP concentrations in control cultures increased over time as biomass accumulated during exponential growth. Over this period, C:N, C:P and N:P stoichiometry remained fairly constant, close to the Redfield ratio. Then, when cultures reached stationary phase, PON and POP concentrations stabilized as nitrate and phosphate became limiting. As consequence, C:N and C:P rapidly increased (Fig. 4). This occur as cells shift fixed carbon to lipid biosynthesis (e.g., Frada et al. 2013). By contrast, during infections after 24hr, POC, PON, and POP concentrations rapid declined. This can be attributed to viral-mediated cell lysis and release of cellular contents to the medium as dissolved forms. This agrees with recently reported important loss of dissolved carbon in *E. huxleyi* during infection (Vincent et al. 2023). Nonetheless, differential loss of POC, PON and POP was detected, which led to a consistent relative enrichment of both C and P relative to N in organic matter during infections (Fig. 4). Carbon enrichment can at least partially be attributed to intracellular C-rich lipid accumulation in infected *E. huxleyi* cells (Rosenwasser et al. 2014; Malitsky et al. 2016). The reason for P enrichment is less clear. The slightly prolonged residual uptake of phosphate detected over the second day of infection, possibly contributed to it, by supply building-blocks for biosynthesis of P-rich macromolecules. Coincidently, lipidome analyses indicated that along marked increments in triacylglycerols in infected cells, there is also a clear buildup in phospholipid pools in infected *E. huxleyi* cells (Malitsky et al. 2016). Furthermore, it is possible that P-rich macromolecules are preferentially retained in decaying cells relative to N-rich macromolecules. Shifts in elemental composition in the “brown tide” chrysophyte *Aureococcus anaphagefferens* has been detected during viral infection (Gobler et al. 1997). Curiously, in that system, N and P were both more retained in the particulate fraction relative to C. It was proposed that cytoplasmic carbon is rapidly lost to the dissolved fraction. By contrast, nitrogen was suggested to be rapidly transferred to concurrent bacteria, while phosphorus may have been preferentially retained in large structural components in algal lysates (Gobler et al. 1997). Each host-viral system likely has its own particularity. However, it is indeed possible that P enrichment in infected *E. huxleyi* resulted from a similar preferential retention in cellular lysates of large structural components, as for example membrane lipid that contain P. Finally, viruses are P-rich particles (Jover et al. 2014). For example the *Paramecium bursaria Chlorella virus*-1 (PBCV-1) that is phylogenetically and size wise close to EhV has been estimated to displays a C:N:P stoichiometry of 17:5:1 (Clasen and Elser 2007; Jover et al. 2014). Thus, the accumulation of viruses inside host cells and associated with cell lysates could produce shifts in organic matter stoichiometry exacerbating P enrichment. Future detailed examination of the elemental and macromolecular redistribution during host cell and viral interplay should enable to decipher the mechanisms underlying stoichiometric shifts during *E. huxleyi* infections following the observations in this study.

In the period post-infection, remaining coccolithophore cell abundance was very low (10^3^–10^4^ cell mL^−1^), while bacterial densities progressively increased to high densities (Fig. 4), most likely reusing organic materials both dissolved or in aggregates typically observed following the viral-mediated lysis of *E. huxleyi* cells (Vincent et al. 2021). Noticeably, over the course of this period, POC, PON and POP concentrations progressively bounced back, but the ratios of C:N and C:P significantly declined. Using a stoichiometric approach (Fig. 5), we estimate such post-infection enrichment in N and P relative to C in organic matter largely resulted from the progressive accumulation of heterotrophic bacteria with typical higher demands in N and P relative to C (C:N= 4.9; C:P=77.0) than phytoplankton (Zimmerman et al. 2014; White et al. 2019). Heterotrophic bacteria triggered by viral-mediated algal lysis, accumulate at high rates in remaining cell-lysates, leading over time to a significant replacement of algal biomass by bacterial biomass in the decaying particulate organic matter.

### 4.3 Potential ecological and biogeochemical implications of EhV infections

During *E. huxleyi* infections, large fractions of organic matter are lost to the dissolved phase as detected by marked POC, PON, and POP concentration decline (Fig. 3). The rapid release of cell contents during infections, as part of the so-called ‘viral shunt’ (Fuhrman 1999; Wilhelm and Suttle 1999), triggers the rapid rise of bacteria. These can actively benefit from the break-down and reuse of dissolved elements and lysates, as it has been shown in other algae-virus systems (e.g., (Gobler et al. 1997). In parallel, stoichiometric variations were detected relative to non-infected cells (Fig. 4). In the short term, during the course of infection and population decline, an enrichment in C and P relative to N was detected, which can be directly attributed to underlying increased metabolic fluxes towards lipid accumulation with *E. huxleyi* virocells, but also differential retention in cell lysates of P-rich molecules. In the longer term, decaying organic matter of *E. huxleyi* cells appears to be progressively replaced to a large extent by bacterial biomass likely bound to aggregates (Vincent et al. 2021) produced during infection.

The current view is that the formation of coccolith-dense aggregates during *E. huxleyi* blooms infected by EhV can enhance the carbon export efficiency, following the ‘viral shuttle’ model (Weinbauer 2004; Sullivan et al. 2017; Laber et al. 2018). However, modifications in the elemental composition can impact the efficiency of carbon export. Rapid export of recently infected populations with high carbon ratios may indeed benefit the efficiency of carbon export. By contrast, slow export or resuspension of lysates in more turbulent settings could result in the enrichment of phosphorus relative to carbon in exported material due to bacterial accumulation. This effect can in turn attenuate the magnitude of the biological carbon pump by lowering the relative contribution of carbon in exported biomass (Finkel et al. 2010b; Deutsch and Weber 2012; Galbraith and Martiny 2015).

Finally, as detailed above, seawater alkalinity draw-down was rapidly stalled during infection, consequently indicating a cessation of cellular calcification. This EhV-mediated effect highlights the antagonistic role of viral infections on the marine carbon counter pump, reducing the net increase of dissolved CO_2_ driven by calcification in growing coccolithophore populations and consequently upholding the buffer capacity of the oceans (Rost and Riebesell 2004).

## 5 Conclusions

In this study, we demonstrate that EhV infection arrests *E. huxleyi* calcification, nutrient acquisition and caused ensuing shifts in the stoichiometry of organic matter. These included both a short-term shift during active viral infection and a long-term shift driven by cohabiting bacterial communities actively consuming algal lysates. During natural blooms, the dynamic of EhV infections can be rapid, but also heterogeneous between different blooming areas (Laber et al. 2018). Thus, one can expect the stoichiometric seascape to be very variable. Nonetheless, such viral-mediated changes in the nature of exported materials, likely influence the cycling of elemental in the photic layer and the export efficiency of biomass to the deep ocean during large-scale coccolithophore blooms. Further research is required to validate our observation in natural coccolithophore populations. However, our results indicate that assessing the impact of viral infections on the elemental biogeochemistry of phytoplankton biomass is crucial to determine their role and impact in the marine ecosystem.

## Data availability

All data generated during the current study are available at the Marine Data Archive (https://mda.vliz.be/directlink.php?fid=VLIZ_00000842_64e4ba6e5d033845577610).

## Supplement

Supplement information contains Table S1 which summarizes the C: N values for heterotrophic bacteria used for the mathematical calcumations presented in Figure 5.

## Author Contributions

T.D., G.A. and M.J.F. conceptualized and designed the research; T.D., G.A., A.P., S.S. and M.J.F. performed research; G.A., A.P. and M.J.F. contributed analytic tools and funding; T.D., G.A., A.P., S.S. and M.J.F. analyzed data and revised paper drafts; T.D. and M.J.F. wrote the paper.

## Competing interests

The authors declare that they have no conflict of interest.

## Acknowledgements

This study was supported by a scholarship from the Bester Award for Oceanography from the Hebrew University of Jerusalem attributed to TD, the Fulbright and Yad Hanadiv Fellowships attributed to SS, the Israel Science Foundation grant number 2921/20 attributed to MJF, the Israel Science Foundation grant number 2361/19 attributed to GA, and by the Natural Sciences and Engineering Research Council of Canada Discovery grant (RGPIN-2022-04305) and Fonds de Recherche du Québec Relève Professorale grant (314248) attributed to AP. We thank Tania Rivlin for undertaking inorganic nutrient measurements and Gil Koplovitz for help during experiments. We finally thank Maoz Fine and Nir Keren for constructive feedback during the preparatory stages of the work.

## Notes

https://mda.vliz.be/directlink.php?fid=VLIZ_00000842_64e4ba6e5d033845577610

### Competing Interest Statement

The authors have declared no competing interest.

### Summary of Updates

Various part of the manuscript were rewritten to improve the description of the results. changes include the title, abstract and sections of the results and discussion in the main text. No novel results were included.

