## Supplemental Table 1 for "Viral infection arrests coccolithophore calcification and nutrient consumption, and triggers shifts in organic stoichiometry"

**Table S1: C: N values for heterotrophic bacteria.**

| **Bacteria species** | **Isolate** | **Average C:N** | **Reference** |
| --- | --- | --- | --- |
| SAR 11 | HTCC1062 | 4.5 | White et al. 2019 |
| SAR 11 | HTCC7211 | 4.5 | White et al. 2019 |
| *Rugeria pomeroy* | DSS-3 | 6.61 | Zimmerman et al. 2014 |
| *Roseobacter (Oceanicola)* | HTCC2516 | 5.12 | Zimmerman et al. 2014 |
| *Roseobacter (Pelagibaca)* | HTCC2601 | 5.56 | Zimmerman et al. 2014 |
| *Alteromonas* | Alt1C | 4.08 | Zimmerman et al. 2014 |
| *Vibrio* | Vib1A | 4.42 | Zimmerman et al. 2014 |
| *Vibrio* | Vib2D | 7.35 | Zimmerman et al. 2014 |
| *Marinomonas* | Oce241 | 4.63 | Zimmerman et al. 2014 |
| *Marinomonas* | Oce340 | 4.6 | Zimmerman et al. 2014 |
| *Psychrobacter* | Mor224 | 4.57 | Zimmerman et al. 2014 |
| *Psychrobacter* | Mor119 | 4.08 | Zimmerman et al. 2014 |
| *Halomonas* | Hal005 | 4.47 | Zimmerman et al. 2014 |
| *Halomonas* | Hal146 | 4.4 | Zimmerman et al. 2014 |
| Total mean | | 4. 92 |  |
